# AtacWorks: A deep convolutional neural network toolkit for epigenomics

**DOI:** 10.1101/829481

**Authors:** Avantika Lal, Zachary D. Chiang, Nikolai Yakovenko, Fabiana M. Duarte, Johnny Israeli, Jason D. Buenrostro

## Abstract

We introduce AtacWorks (https://github.com/clara-genomics/AtacWorks), a method to denoise and identify accessible chromatin regions from low-coverage or low-quality ATAC-seq data. AtacWorks uses a deep neural network to learn a mapping between noisy ATAC-seq data and corresponding higher-coverage or higher-quality data. To demonstrate the utility of AtacWorks, we train a model on data from four human blood cell types and show that this model accurately denoises chromatin accessibility at base-pair resolution and identifies peaks from low-coverage bulk sequencing of unseen cell types and experimental conditions. We use the same framework to obtain high-quality results from as few as 50 aggregate single-cell ATAC-seq profiles, and also from data with a low signal-to-noise ratio. We further show that AtacWorks can be adapted for cross-modality prediction of transcription factor footprints and ChIP-seq peaks from low input ATAC-seq. Finally, we demonstrate the applications of our approach to single-cell genomics by using AtacWorks to identify regulatory regions that are differentially-accessible between rare lineage-primed subpopulations of hematopoietic stem cells.

## Introduction

Within a single cell, the eukaryotic genome is hierarchically organized to form a gradient of chromatin accessibility ranging from compact, repressive heterochromatin to nucleosome-free regions associated with increased gene expression. Assay for Transposase-Accessible Chromatin using Sequencing (ATAC-seq) leverages the Tn5 transposase to directly measure chromatin accessibility as a proxy for the relative activity of DNA regulatory regions across the genome^1^. ATAC-seq has been applied to identify the effects of transcription factors on chromatin, construct cellular regulatory networks, and localize epigenetic changes underlying diverse development and disease-associated transitions^2–4^. Recently, the development of single-cell ATAC-seq methods have made it possible to measure accessible chromatin in individual cells, enabling epigenomic analysis of rare cell types within heterogeneous tissues^5^.

The ability to measure biologically-meaningful changes in accessible chromatin using ATAC-seq depends on both the signal-to-noise ratio and the depth of sequencing coverage. Technical parameters such as the overall quality of cells or tissues, the nuclei extraction method^6^, or over-digestion of chromatin can result in attenuated measurements of accessibility. Importantly, these issues are exacerbated in single-cell experiments, where primary tissues may vary in quality and key cell types may be exceedingly rare.

Deep learning represents a potential tool to address these limitations, as it has been successfully used for problems such as denoising speech^7^ and image restoration^8,9^. An earlier study demonstrated that simple convolutional neural networks can be used to denoise and call peaks from ChIP-seq data, but was optimized for broad peak calling of histone modifications^10^. Another recent study applied deep learning to predict chromatin accessibility in a rare pancreatic islet cell type^11^, highlighting the need for a robust and generalizable method for the analysis of sparse ATAC-seq data.

Here, we introduce AtacWorks, a deep learning-based toolkit that takes as input a low-coverage or low-quality ATAC-seq signal, and denoises it to produce a higher-resolution or higher-quality signal. AtacWorks trains a model to accurately predict both chromatin accessibility at base-pair resolution (a coverage track), and the genomic locations of accessible regulatory regions (peak calls). We apply AtacWorks to subsampled low-coverage bulk ATAC-seq and show that AtacWorks improves the resolution of the chromatin accessibility signal and the identification of regulatory elements. Further, AtacWorks is able to denoise signal from cell types not present in the training set, demonstrating that our deep learning models learn generalizable features of chromatin accessibility. We use the same framework to denoise aggregated single-cell ATAC-seq from a small number of cells, and also to improve the signal-to-noise ratio in an ATAC-seq dataset with low signal-to-noise. We further show that AtacWorks can be adapted for cross-modality prediction of transcription factor footprints and ChIP-seq peaks from low input ATAC-seq. Finally, we apply AtacWorks to single-cell ATAC-seq of hematopoietic stem cells to identify regulatory elements associated with rare lineage-primed subpopulations.

## Results

### A deep learning framework for denoising low-coverage data

AtacWorks trains a deep neural network to learn a mapping between noisy, low-coverage or low-quality ATAC-seq data and matching high-coverage or high-quality ATAC-seq data from the same cell type. Given a noisy ATAC-seq signal track as input, a trained model performs two tasks: denoising at base-pair resolution (predicting an improved signal track) and peak calling (predicting the genomic location of accessible regulatory elements). Once this mapping is learned, it is saved as a model that can be applied to denoise and call peaks from similar low-coverage or low-quality datasets at any given region in the genome.

The network makes predictions for each base in the genome based on coverage values from a surrounding region spanning several kilobases (6 kb for the models presented here), but does not consider the DNA sequence itself, allowing it to generalize across cell types. AtacWorks uses the Resnet (residual neural network) architecture, which has been applied extensively for natural image classification and localization^12^. Our architecture consists of multiple stacked residual blocks, each composed of three convolutional layers and a skip connection that bypasses intermediate layers (Fig. 1a). These skip connections allow propagation of the input through the layers of the network to avoid vanishing gradients^12^, enabling deeper and more accurate models to be trained. The model is trained using a multi-part loss function combining Mean Squared Error (MSE), 1 - Pearson Correlation, and Binary Cross-Entropy (BCE) losses (see Methods).

**Figure 1.**
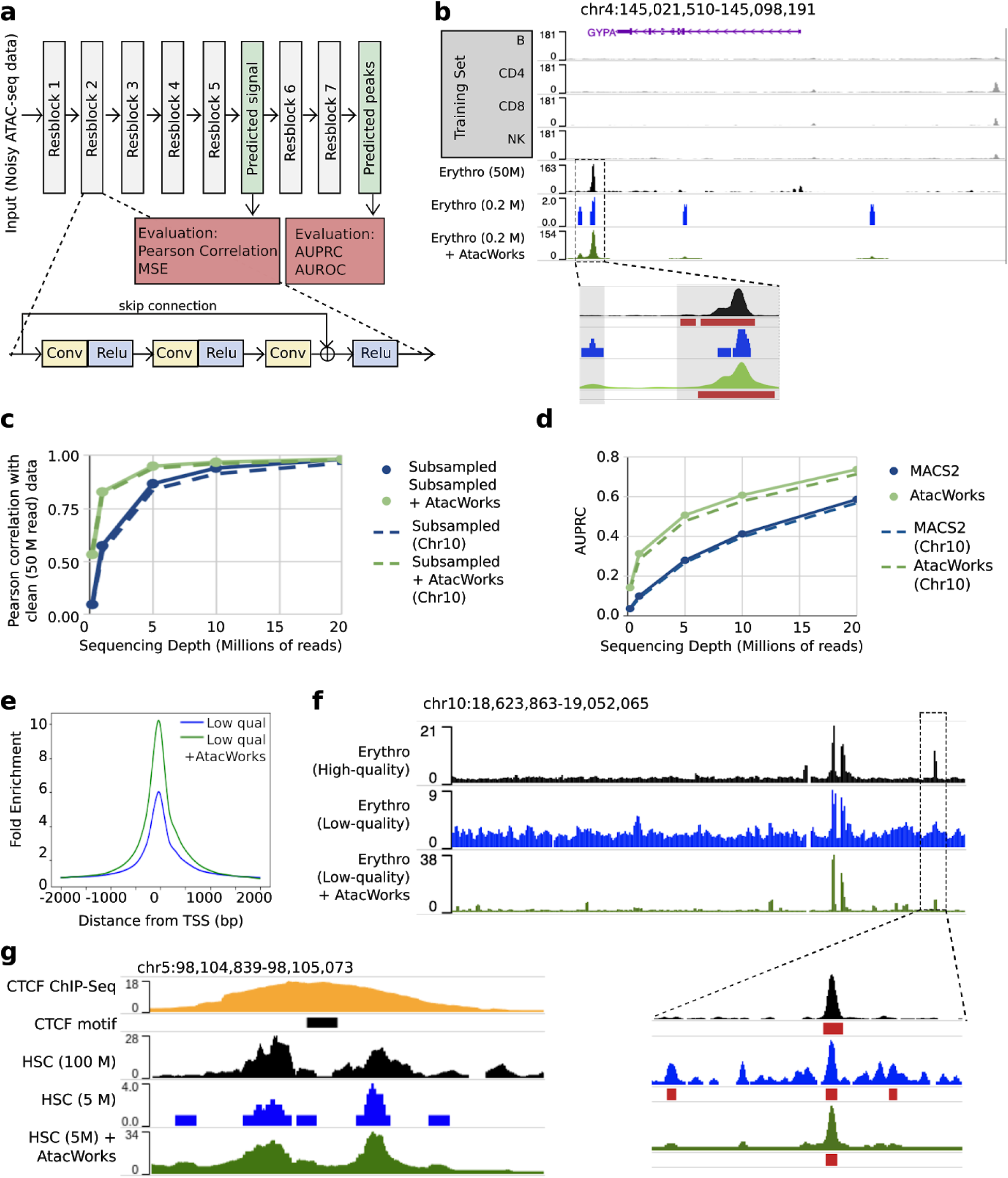
A deep learning approach to denoise ATAC-seq data. **A.** Schematic of the Resnet architecture. The zoomed-in region displays a residual block composed of 1-dimensional convolutional layers (Conv), nonlinear activation functions (ReLU), and a skip connection. **B**. ATAC-seq signal tracks near the erythroblast marker gene *GYPA*, for four cell types used to train an AtacWorks model (gray), high-coverage erythroblast data (50 million reads; black), and erythroblast data subsampled to 0.2 million reads before (blue) and after (green) denoising with AtacWorks. Red bars below the zoomed-in tracks show peak calls by MACS2 (for the 50 M and 0.2 M read tracks) and AtacWorks (for the denoised track). **C**. Pearson correlation between a clean ATAC-seq signal track (50 million reads) and subsampled data for erythroblasts, before (blue) and after (green) denoising with AtacWorks. Solid lines show correlation over the genome; dotted lines show correlation over chromosome 10. **D**. AUPRC for MACS2 (blue) and AtacWorks (green) showing their peak calling performance on subsampled data, using peaks called by MACS2 subcommands on the clean (50 million reads) signal track as truth. Whole lines show AUPRC over the genome; dotted lines show AUPRC over chromosome 10. **E**. Average ATAC-seq signal over 4000-bp windows centered on transcription start sites, in low-quality ATAC-seq data from erythroblasts, before (blue) and after (green) denoising with AtacWorks. **F**. ATAC-seq signal tracks for high-quality (black) and low-quality (blue) data from erythroblasts, and low-quality data after denoising by AtacWorks (green). Red bars below the tracks show peak calls by MACS2 (for the high and low-quality tracks) and AtacWorks (for the denoised track). **G.** Signal tracks around a CTCF binding site in HSCs. From top to bottom: CTCF ChIP-seq signal (yellow), ATAC-seq signal at a depth of 100 million reads (black), subsampled to 5 million reads (blue), and subsampled signal denoised by AtacWorks (green). The black bar underneath the ChIP-seq track shows the CTCF binding motif.

We used AtacWorks to train deep learning models with bulk ATAC-seq data from FACS-isolated human blood-derived cell types^2^. To do this, we obtained ATAC-seq datasets from 4 cell types (B cells, NK cells, CD4^+^ and CD8^+^ T cells) and sampled each to a depth of 50 million reads (25 million read pairs) to produce standardized clean (high-coverage) data. Peaks for each clean dataset were identified using MACS2 (see Methods). We then subsampled each clean dataset to multiple lower sequencing depths ranging from 0.2 million to 20 million reads (Supplementary Fig. 1). For each depth, we trained a model to take as input the low-coverage ATAC-seq signal and reconstruct both the clean ATAC-seq signal and peak calls.

To assess the generalizability of our method, we tested the performance of these models on ATAC-seq data from erythroblasts^2^, which were not included in the training set. We first subsampled reads from erythroblasts to the same depths as the training data. For each sequencing depth, we then applied the trained model to the corresponding subsampled dataset to obtain a predicted high-coverage signal track and peak calls. By examining the resulting denoised tracks, we confirmed that AtacWorks identifies cell-type-specific peaks that were not present in the training data, including a region adjacent to erythroblast marker gene *GYPA*^*2*^ (Fig. 1b). This suggests that our models are learning generalizable features of chromatin accessibility rather than cell-type specific patterns.

To quantitatively evaluate the denoised high-coverage signal tracks produced by AtacWorks, we compared them to a clean (50 million read) erythroblast signal. At all sequencing depths, the Pearson correlation, Spearman correlation, and MSE between the denoised and clean signal tracks were substantially greater than that between the noisy and clean signal, both within and outside accessible chromatin peaks (Fig. 1c, Supplementary Table 1, Supplementary Fig. 2). We further found that our method outperforms smoothing using linear regression using these metrics (Supplementary Table 2). Next, we evaluated the peaks identified by AtacWorks from each sequencing depth, and found that both the Area Under the Precision-Recall Curve (AUPRC) and Area Under the Receiver-Operator Characteristic (AUROC) of peaks were superior to MACS2^13^ called peaks from the same subsampled data (Fig. 1d, Supplementary Table 1, Supplementary Fig. 2). For this analysis, AtacWorks produced output data of quality equivalent to (on average) 2.6x the number of reads in the input data based on Pearson correlation, and 4.2x based on AUPRC (Supplementary Table 1).

**Figure 2.**
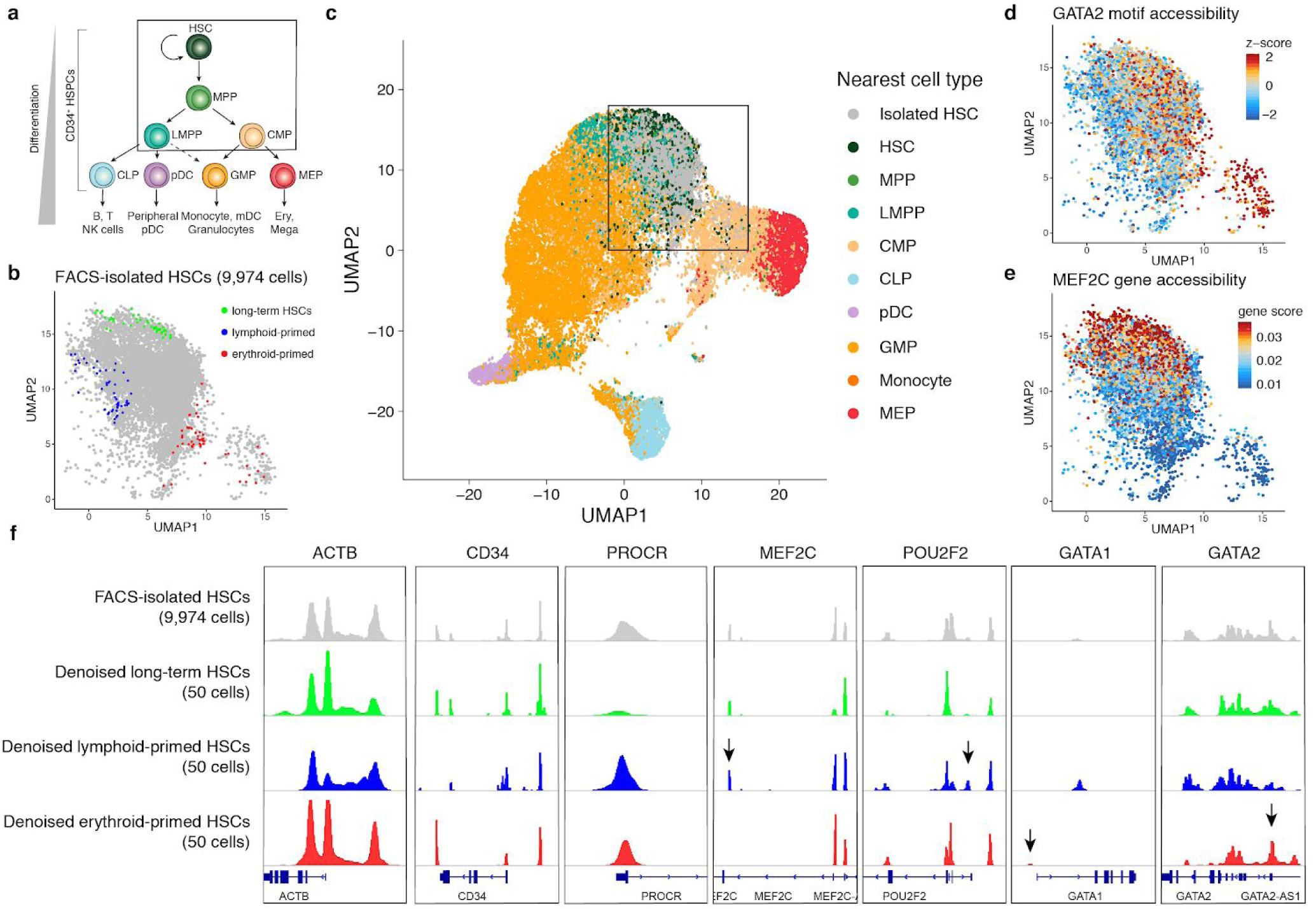
AtacWorks identifies differentially-accessible regulatory regions associated with lineage-primed hematopoietic stem cells. **A.** A schematic of the classical hierarchy of human hematopoietic differentiation. **B.** A UMAP dimensionality reduction of single-cell ATAC-seq profiles from 9,974 FACS-isolated hematopoietic stem cells (HSCs). The colored points represent three 50-cell subsamples, each generated by selecting a single cell and identifying its nearest neighbors in principal component space. **C.** A combined UMAP dimensionality reduction of single-cell ATAC-seq profiles from HSCs shown in (B) and 28,505 previously-published bead-enriched CD34^+^ bone marrow progenitor cells^14^. The bead-enriched CD34^+^ cells are colored by the most correlated cell type from a FACS-isolated single-cell ATAC-seq reference^18^. The box indicates the region containing FACS-isolated HSCs shown in (B), (D) and (E). **D.** FACS-isolated HSCs colored by chromVAR transcription factor motif accessibility z-scores (see Methods) for GATA2. **E.** FACS-isolated HSCs colored by smoothed gene accessibility scores (see Methods) for *MEF2C*. **F.** Aggregate chromatin accessibility signal tracks surrounding genes implicated as markers of lineage priming^18,23^ for all 9,974 FACS-isolated HSCs and the three denoised 50-cell subsamples of HSCs shown in (B). The arrows denote select regulatory regions with significant differences in chromatin accessibility relative to a permuted background.

To show that the models are not simply learning features specific to the training set, we calculated performance metrics on chromosome 10, which was previously held-out from training, and obtained highly similar results to those computed on the whole genome (Fig. 1c and 1d, Supplementary Table 1). We also evaluated model performance specifically on differential peaks present in only either the training or test set, and found that AtacWorks improves both the signal track accuracy and peak calling in these regions (Supplementary Table 1). Further, we found that the results were highly robust to different subsets of the training data used (Supplementary Table 3). We obtained similar results by applying blood cell-trained models to subsampled low-coverage ATAC-seq data from human cells derived from the Peyer’s Patch, suggesting our models are robust to experimental preparation of cells (Supplementary Fig. 3, Supplementary Table 4).

Finally, we compared our method to a recent study^11^ that also reported the use of a deep learning model that could perform either ATAC-seq denoising or peak calling. We implemented the U-Net model architecture reported in this study and found that the Resnet architecture used in AtacWorks outperforms this model in denoising, peak calling, and runtime (Supplementary Note 1, Supplementary Table 5).

### AtacWorks generalizes to diverse applications

To demonstrate our method is also broadly adaptable to bleeding edge use cases of ATAC-seq, we applied AtacWorks to denoise data from a high-throughput single-cell ATAC-seq experiment. We first obtained droplet single-cell ATAC-seq (dscATAC-seq) data from bead-isolated human blood cells and aggregated single-cell chromatin accessibility profiles by cell type^14^. We then trained AtacWorks models on data from two cell types (B cells and monocytes), randomly sampling 10 cells (∼0.2 million reads) or 50 cells (∼1 million reads) for the low-coverage training datasets. The resulting trained models improved signal track accuracy and peak calling from aggregated natural killer (NK) cells sequenced using the same protocol (Supplementary Table 6). AtacWorks improved the AUPRC of peak calls from 50 NK cells from 0.2048 to 0.7008, a result that MACS2 requires over 400 cells to obtain (Supplementary Table 6). We also tested AtacWorks on monocytes sequenced using a combinatorial indexing-based variant of the same protocol (dsciATAC-seq) and saw similar improvements (Supplementary Table 7, Supplementary Fig. 4).

Another present challenge in adapting ATAC-seq to novel biological contexts is developing experimental protocols that optimize enrichment of open chromatin. To help address this issue, we applied AtacWorks to improve signal quality in ATAC-seq datasets with low signal-to-noise ratio. We trained a model to learn a mapping between paired high and low-quality ATAC-seq datasets from FACS-isolated human monocytes^2^ (Supplementary Table 8, Methods). Both datasets had similar sequencing depth (approximately 20 million reads); however, one had a higher signal-to-noise ratio estimated using the global enrichment of signal surrounding transcription start sites (TSSs). We then applied this trained model to denoise low-quality bulk ATAC-seq data of similar depth from erythroid cells. AtacWorks improved the enrichment at TSSs (Fig. 1e), producing a signal track and peak calls more similar to those obtained from higher-quality data (Fig. 1f, Supplementary Table 9).

### AtacWorks enables cross-modality predictions

Seeing that AtacWorks accurately predicts denoised coverage at base-pair resolution, we sought to extend it for transcription factor footprinting^1,15^. “Footprinting” leverages the fact that transcription factors vary in how they bind to DNA, which allows binding events to be identified via a characteristic Tn5 insertion signature. Traditionally, footprinting requires over 100 million reads^15^, prohibiting its widespread use. To test the feasibility of performing footprinting from low-input ATAC-seq, we obtained high-coverage (100 million reads) ATAC-seq data from FACS-sorted human blood cell types (MPP cells, CD8+ T cells, NK cells)^2^ and reduced track smoothing to preserve transcription factor-specific patterns of Tn5 insertions (see Methods). We then downsampled these tracks to lower sequencing depths and trained a model for each depth, which we tested on data from similarly-processed HSCs. We evaluated the performance of these models on a set of 200-bp genomic regions spanning binding motifs for genome architectural protein CTCF. At all sequencing depths, AtacWorks improved the signal track spanning CTCF motifs in HSCs (Supplementary Table 10), enhancing the characteristic footprint of CTCF binding (Fig. 1g, Supplementary Fig. 5).

Encouraged by these results, we reasoned we may adapt our method to directly predict ChIP-seq peaks from low-input ATAC-seq. Like footprinting, standard ChIP-seq protocols also require large quantities of input material (at least 10^7^ cells), though this number has been reduced in certain contexts by recent developments^16^. To demonstrate the feasibility of cross-modality prediction, we trained AtacWorks models to learn a mapping from low-coverage aggregate dscATAC-seq signal to CTCF and H3K27ac (an active histone mark) ChIP-seq signal and peak calls in the same cell type. For the prediction of CTCF ChIP-seq, we also supplied the model with the positions of CTCF binding motifs on both strands of the genome (see Methods). We trained models on noisy aggregate dscATAC-seq data from small numbers of B cells, and tested them on similarly-processed monocytes. For small numbers of cells ranging from 10 to 500, AtacWorks predicted CTCF and H3K27ac peak calls with surprisingly high concordance to ChIP-seq data from the same cell type (AUROC 0.9 from 500 cells; Supplementary Fig. 6, Supplementary Table 11). Altogether, these results demonstrate the potential for AtacWorks to be broadly applied for cross-modality inference of latent epigenetic states.

### AtacWorks enhances the resolution of single-cell studies

Empowered by the improved resolution afforded by AtacWorks, we sought to investigate epigenetic changes underlying development in rare subpopulations of cells that cannot be experimentally isolated, and thus cannot be analyzed using traditional approaches. Previous single-cell studies of FACS-isolated bone marrow mononuclear cells (BMMCs) have observed epigenetic heterogeneity within immunophenotypically-defined cellular populations, suggesting that hematopoietic stem and progenitor cells lie along a continuum of differentiation states (Fig. 2a)^17,18^. In particular, hematopoietic stem cells (HSCs) are thought to include rare subpopulations of cells that are primed towards either the lymphoid or erythroid lineage^17,19,20^. Though single-cell ATAC-seq enables measurements of chromatin accessibility over aggregate genomic features, such as sets of transcription factor motifs^21^ or the regions surrounding transcription start sites^21^,22, with such granular lineage-primed states, there is typically not enough sequencing coverage to identify which specific regulatory regions are associated with each differentiation trajectory.

We reasoned we could use AtacWorks to identify sets of regulatory regions that are unique to lymphoid or erythroid-primed HSCs. First, we performed dscATAC-seq^14^ on FACS-isolated HSCs to generate 9,974 single-cell chromatin accessibility profiles (see Methods). To define lymphoid and erythroid differentiation trajectories, we collected published dscATAC-seq data from bead-enriched CD34^+^ cells and used a bulk reference-guided approach (see Methods) to project all single-cell profiles into a shared latent space, visualized using UMAP for dimensionality reduction (Fig. 2b, c). This analysis localized FACS-isolated HSCs to a region at the top of the projection. We then confirmed that HSCs localized in this region exhibited directional signal bias in transcription factor motif accessibility scores for the GATA2 motif (Fig. 2d) and smoothed gene accessibility scores for *MEF2C* (Fig. 2e), genes which have been implicated as markers of erythroid and lymphoid lineage priming respectively^18,23^ (see Methods).

To generate high-resolution chromatin accessibility tracks of lineage-primed cells using our model, we selected three distant samples of 50 HSCs each, representing putative populations of long-term, lymphoid and erythroid-primed HSCs (Fig. 2b). For each sample of 50 aggregated cells, we performed signal denoising using AtacWorks and visualized the denoised chromatin accessibility profiles near genes suggested to be indicators of lineage priming^18,23^ (Fig. 2f). We observed considerable differences between the denoised tracks that could not be readily distinguished from the original low-coverage signal (Supplementary Fig. 7), including potential regulatory elements seemingly present in the lymphoid, but not the erythroid-primed cells (near *MEF2C, POU2F2*) and vice-versa (near *GATA1, GATA2*).

To access the significance of these chromatin accessibility differences, we took 1000 samples of 50 randomly-selected HSCs and used AtacWorks to denoise a list of select genomic regions (200 kb windows surrounding 2,303 genes with differential expression across CD34^+^ cells, see Methods). For each of the ∼300 million bases in the selected regions, we calculated a normalized mean and standard deviation of the coverage from the 1000 denoised tracks, allowing us to estimate z-scores for each regulatory region we observed in our denoised long-term HSC and lineage-primed samples (see Methods). We identified an average of 2,839 regulatory regions with significant differential accessibility per sample (Supplementary Table 12), including those highlighted (Fig. 2f). Though these regulatory elements have yet to be functionally validated, these results demonstrate the unique capacity of deep learning to enhance the resolution of sparse single-cell ATAC-seq studies.

## Discussion

ATAC-seq has become a widely adopted tool for high-resolution characterization of the epigenome, providing insights into the mechanisms underlying gene expression changes associated with development, evolution, and disease. However, technical limitations in tissue quality, assay performance, and sequencing coverage constrain our ability to measure the full spectrum of chromatin states across the genome. These limitations also pertain to emerging single-cell ATAC-seq technologies, as cell types of interest are often difficult to experimentally isolate, and are present at low frequencies in heterogeneous contexts.

Here we present AtacWorks, an easy-to-use and generalizable toolkit to train and apply deep learning models to ATAC-seq data. Unlike previous deep learning methods for epigenomics, AtacWorks denoises ATAC-seq signal at base-pair resolution and simultaneously predicts the genomic location of accessible regulatory elements. The models we present here outperform existing approaches at both of these tasks, and moreover, are robust across cell and tissue types, individuals, and experimental protocols. We also demonstrate that AtacWorks can be adapted for cross-modality prediction of transcription factor footprints and ChIP-seq peaks from low-input ATAC-seq. As such, we anticipate this framework may be broadly useful for other deep learning applications in genomics, such as DNase, MNase, ChIP-seq, and the recently-developed method CUT&RUN^16^.

Finally, the robustness and speed of AtacWorks enable its application to high-throughput single-cell ATAC-seq datasets of heterogeneous tissues. We show that our method can be used on small subsets of rare lineage-priming cells to denoise signal and identify accessible regulatory regions at previously-unattainable genomic resolution. Based on these advancements, we anticipate that AtacWorks will broadly enhance the utility of epigenomic assays, providing a powerful platform to investigate the regulatory circuits that underlie cellular heterogeneity.

## Methods

### Data preprocessing

BAM files for bulk ATAC-seq were downsampled to a fixed number of reads using SAMtools v.1.9^24^. For paired-end sequencing data, both reads in a read pair were retained (e.g. 100,000 read pairs were selected to obtain a total of 200,000 sequencing reads). For CTCF footprinting experiments, downsampling was repeated independently 5 times to produce 5 times the amount of training data. This was done to ensure that the model received enough training data, as only a small fraction of the genome was used for training in these experiments.

For single-cell ATAC-seq experiments, a number of cells of the chosen cell type were randomly selected and all reads from those cells were extracted from the BAM file via cell barcodes. This random sampling of cells was repeated independently 5 times due to the sparsity of the input single-cell ATAC-seq data.

To identify the exact location of Tn5 insertions with base pair resolution, each ATAC-seq read was converted to a single genomic position corresponding to the first base pair of the read. Previous work has demonstrated that the Tn5 transposase inserts adapters separated by 9 bp, so reads aligning to the + strand were offset by +4 bp, while reads aligning to the - strand were offset by -5 bp^1^. Each cut site location was extended by 100 bp in either direction, except for transcription factor footprinting experiments where each cut site was extended by 5 bp in either direction. The bedtools genomecov function^25^ was then used to convert the list of locations into a genome coverage track containing the ATAC-seq signal at each genomic position.

To call peaks from clean and noisy signal tracks, MACS2 subcommands bdgcmp and bdgpeakcall were run with the ppois parameter and a -log10(p-value) cutoff of 3. BED files with equal coverage over all chromosomes were provided as a control input track.

### Running AtacWorks

#### Input data

Deep learning models were trained using one or more pairs of matching ATAC-seq datasets. Each pair consisted of two ATAC-seq datasets from the same sample or cell type: a “clean” dataset of high sequencing coverage or quality, and a “noisy” dataset of lower coverage or quality. Unless indicated otherwise, low-coverage datasets were generated computationally by randomly subsampling a fraction of reads or cells from the high-coverage dataset.

Models were given three inputs for each such pair of datasets:

1. a signal track representing the number of sequencing reads mapped to each position on the genome in the noisy dataset.
2. a signal track representing the number of sequencing reads mapped to each position on the genome in the clean dataset.
3. The genomic positions of peaks called by MACS2 on the clean dataset.

Models learned a mapping from (1) to both (2) and (3); in other words, from the noisy signal track, they learned to predict both the clean signal track, and the positions of peaks in the clean dataset.

#### Dividing data into genomic intervals

Input ATAC-seq datasets were divided into training, validation and holdout sets. The validation set consisted of data for chromosome 20, while the holdout set consisted of data for chromosome 10. Datasets for all other autosomes were included in the training set. These datasets were then further divided into non-overlapping intervals of 50 kb, unless otherwise specified (Supplementary Table 13), each representing a single training example. Each 50-kb long interval was padded with an additional 5 kb at either end, unless otherwise specified (Supplementary Table 13) so that the convolutional filter had enough neighboring bases to make predictions for every base inside the interval. Balanced training datasets with a fixed proportion of peaks were tested; however, this feature did not improve the overall performance metrics and was therefore not employed in genome-wide experiments.

#### Resnet architecture

The PyTorch neural network framework^26^ was used to train a Resnet (residual neural network) model consisting of multiple stacked residual blocks. Each residual block included three convolutional layers and a skip layer to add the input to the first layer to the output of the third layer (Fig. 1a). Unless specified otherwise (Supplementary Table 14), each convolutional layer used 15 convolutional filters with a kernel size of 51 and a dilation of 8. Dilated convolutional layers were used to increase the receptive field of the model without increasing the parameter count. This approach has been effective in image classification tasks where a larger receptive field is desirable^27^. Models did not utilize batch normalization^28^ for the convolutional layers, as it did not improve accuracy on either the regression or classification tasks in our experiments.

For each position in the given interval, the model performed two tasks; a regression or denoising task (predicting the ATAC-seq signal at each position) and a classification or peak calling task (predicting the likelihood that each position is part of a peak).

In order to perform both tasks, the input was passed through several residual blocks, followed by a regression output layer that returns the predicted ATAC-seq signal at each position in the input. The regression output was then passed through another series of residual blocks followed by a classification output layer that returned a prediction for whether each base in the input is part of a peak.

The rectified linear unit (ReLU) activation function was used throughout the network, except for the classification output layer, which used a sigmoid activation function. The sigmoid activation forced the network to return a value between 0 and 1 for each input base, which was interpreted as the probability of that base being part of a peak. A cutoff of 0.5 was used to call peaks from these probability values.

Other convolutional neural network architectures, including the U-Net^29^ were tested, and the selected architecture was chosen based on its robust performance in both denoising and peak calling tasks on several datasets.

#### Model training

All deep learning models were trained using a multi-part loss function, comprising a weighted sum of three individual loss functions:

1. Mean squared error (MSE; for the regression output)
2. 1 - Pearson correlation coefficient (for the regression output)
3. Binary cross-entropy (for the classification output)

The relative importance of these loss functions was tuned by assigning different weights to each (Supplementary Table 13).

Training examples were randomly shuffled at the beginning of each training epoch and passed to the deep learning model in batches of 64 examples each, unless otherwise specified (Supplementary Table 13). At the end of each epoch of training, the performance of the model on the validation set was evaluated, and the model with the best validation set performance was saved and used.

Models were trained using the Adam optimizer^30^ with a learning rate of 2 x 10^−4^ for 25 epochs.

#### Model evaluation

The performance of the model in regression was measured by computing Pearson correlation, Spearman correlation and MSE of the denoised data with respect to the clean dataset. For classification (peak calling), the model outputs the probability of belonging to a peak, for each position in the genome. In order to obtain predicted peaks, there is a set probability threshold above which a base is said to be a peak. Similarly, MACS2 produces a p-value for each position and the final peak calls depend on a user-defined probability threshold. Therefore the Area under the Precision-Recall Curve (AUPRC) and Area under the Receiver Operating Characteristic (AUROC) metrics were used to evaluate classification performance over the entire range of possible thresholds.

#### Parameters

All the parameters describing the models used in this paper are given in Supplementary Table 14. These parameters were chosen in a grid search based on validation set performance. Deeper and larger models produced slightly better results; however, larger models were also expensive and time-consuming to train.

#### Runtime

AtacWorks took 2.7 minutes per epoch to train on one ATAC-seq dataset, and 22 minutes to test on a different whole genome, using 8 Tesla V100 16GB GPUs in an NVIDIA DGX-1 server.

### Single-cell ATAC-seq denoising experiments

Two experiments were performed to validate the denoising and peak calling performance of AtacWorks on single-cell ATAC-seq data.

The first experiment used dscATAC-seq data from B cells, monocytes, and natural killer cells^14^ (Supplementary Table 6). 2400 cells (∼48 million reads) of each type were randomly selected to generate “clean” high coverage signal tracks and peak calls, and then 10 and 50 cells from among the 2400 cells of each type were subsampled to obtain noisy low-coverage data. The data from B cells and monocytes was used to train the model, which was then tested on data from the NK cells.

The second experiment used dsciATAC-seq data from CD4^+^ T cells, CD8^+^ T cells, pre-B cells, and monocytes^14^ (Supplementary Table 7). 6000 cells (∼13 million reads) of each type were randomly selected to obtain “clean” signal tracks and peak calls, and 90 (∼200,000 reads) or 450 (∼1 million reads) cells of each type were subsampled to obtain noisy data. 3 cell types (CD4^+^ T cells, CD8^+^ T cells, and pre-B cells) were used for training deep learning models. These models were tested on noisy data of the fourth cell type (monocytes).

### Paired high and low-quality tracks

Paired high and low-quality chromatin accessibility tracks were computationally generated from the same experiment in order to minimize the impact of potential batch effects. Previously published bulk ATAC-seq tracks from monocytes and erythroblasts^2^ were split by technical and biological replicate, and then quantified using a TSS enrichment score. Tracks were then visually classified as high or low enrichment, and then aggregated based on classification and cell type to form the paired high and low-quality tracks (Supplementary Table 8). The original study describing these datasets found that ATAC-seq profiles were highly reproducible across both technical and biological replicates (mean Pearson r = 0.94 and r = 0.91, respectively)^2^.

### Adding binding motif locations for CTCF ChIP-seq prediction

The deep learning model was modified to take additional inputs along with the noisy ATAC-seq signal. Potential CTCF (CCCTC-binding factor) binding sites were identified on both strands of the genome using motifmatchr (https://github.com/GreenleafLab/motifmatchr). The top 200,000 sites were selected and expanded to 500 bp regions centered on the known binding motif. In order to predict CTCF ChIP-seq peaks from ATAC-seq data, the model was given the positions of CTCF binding motifs on the genome in addition to the noisy ATAC-seq coverage track. For every position in the genome, the model received three numeric inputs: the coverage at that position in the noisy ATAC-seq dataset, a 0 or 1 representing whether that position was part of a CTCF binding motif on the forward strand, and a 0 or 1 representing whether that position was part of a CTCF binding motif on the reverse strand.

### Applying AtacWorks to lineage-priming hematopoietic stem cells

#### Generation of droplet single-cell ATAC-seq data for FACS-isolated HSCs

Cryopreserved human bone marrow mononuclear cells were purchased from Allcells (catalog number BM, CR, MNC, 10M). Cells were quickly thawed in a 37°C water bath, rinsed with culture medium (RPMI 1640 medium supplemented with 15% FBS) and then treated with 0.2 U/μL DNase I (Thermo Fisher Scientific) in 2 mL of culture medium at room temperature for 15 min. After DNase I treatment, cells were filtered with a 40 μm cell strainer, washed with MACS buffer (1x PBS, 2 mM EDTA and 0.5% BSA), and cell viability and concentration were measured with trypan blue on the TC20 Automated Cell Counter (Bio-Rad). Cell viability was greater than 90% for all samples. CD34 positive cells were then bead enriched using the CD34 MicroBead Kit UltraPure (Miltenyi Biotec, catalog number 130-100-453) following manufacturer’s instructions. The enriched population was then simultaneously stained with CD45, Lineage cocktail, CD34, CD38, CD45RA and CD90 antibodies in MACS buffer for 20 min at 4°C. Stained cells were then washed with MACS buffer and the CD45+ Lin-CD38-CD34+ CD45RA-CD90+ fraction (HSCs) was FACS sorted using a MoFlo Astrios Cell Sorter (Beckman Coulter). Single-cell ATAC-seq data was then generated for the sorted HSCs using the dscATAC-seq Whole Cell protocol as previously described^14^.

#### Preprocessing of dscATAC-seq data

Per-read bead barcodes were parsed and trimmed using UMI-tools ^31^. Constitutive elements of the bead barcodes were assigned to the closest known sequence allowing for up to 1 mismatch per 6-mer or 7-mer (mean >99% parsing efficiency across experiments). Paired-end reads were aligned to hg19 using BWA^32^ on the Illumina BaseSpace online application. Bead-based ATAC-seq processing (BAP)^14^ was used to identify systematic biases (i.e. reads aligning to an inordinately large number of barcodes) and barcode-aware deduplicate reads, as well as perform merging of multiple bead barcode instances associated with the same cell. Barcode merging was necessary due to the nature of the BioRad SureCell scATAC-seq procedure used in this study, which enables multiple beads per droplet. BAP was given an alignment (.bam) file for a given experiment with a bead barcode identifier indicated by a SAM tag as input.

#### Bulk-guided projection

The bulk-guided UMAP projection of single cells (Fig. 2c) was performed as previously described^14^. In brief, a common set of peaks (*k* = 156,311) was used to create a vector of read counts for each CD34^+^ single-cell ATAC-seq profile. Principal Component Analysis (PCA) was run on previously-published bulk ATAC-seq data^2^ to generate eigenvectors capturing variations in regulatory element accessibility across cell types. Each single cell was then projected in the same space as these eigenvectors by multiplying their counts vector by the common PCA loading coefficients. The resulting projection scores were scaled and centered prior to being visualized using UMAP. Predicted labels for the CD34^+^ cells were derived by correlating their projected single-cell scores with those of a reference set of FACS-isolated PBMCs ^18^ and assigning the label of the closest match.

#### Transcription factor motif accessibility z-scores

Motif accessibility z-scores for GATA2 (Fig. 2d) were computed using chromVAR^21^. The method calculates gain or loss in accessibility within peaks that share a common transcription factor motif while adjusting for GC content and overall region accessibility. The single cells were scored using a provided file called “human_pwms_v1” that is a list of human transcription factor motifs curated from the CIS-BP database.

#### Smoothed gene accessibility scores

Gene accessibility scores for MEF2C (Fig. 2e) were computed as previously described^14^. Briefly, to obtain gene scores for a particular gene across all cells, any sequencing reads within 10kb of the gene’s transcription start site were compiled and weighted using an inverse exponential decay function. The weighted reads were then summed for each cell and smoothed by averaging the scores from each cell’s 50 nearest neighbors in principal component space. A list of transcription start sites for hg19 was obtained from the UCSC Genome Browser.

#### Denoising lineage-priming hematopoietic stem cells

Each subsample of lineage-priming hematopoietic stem cells (HSCs) was generated by selecting a single HSC and aggregating the 50 most similar HSCs in principal component space. The selected HSCs were chosen and annotated based on their proximity to specific populations of labeled CD34^+^ cells (Fig. 2b). After aggregation, three resulting subsamples were converted from BAM to bigWig format as described earlier and denoised using a model trained on dscATAC-seq data from B cells and monocytes. The denoised tracks were then normalized by coverage for cross-sample comparisons.

#### Random permutation and denoising of hematopoiesis regulatory regions

A list of differentially expressed genes in blood cells was obtained from the Human Cell Atlas Data Portal and filtered down to a set of 2,303 genes relevant to HSCs. Transcription start sites for each of these genes were obtained and expanded by 100kb in both directions to generate a set of hematopoiesis regulatory regions comprising around 300 million bases, or 10% of the genome.

To provide a background model for the denoised lineage-primed samples, 1000 subsamples containing 50 HSCs each were generated by randomly sampling from the pool of 9,974 HSCs with replacement. These random samples were converted from BAM to bigWig format as described earlier. The 1000 random samples were then denoised using the same AtacWorks model used to denoise the lineage-priming HSCs, but only in defined hematopoiesis regulatory regions, reducing the runtime by over 90%. The denoised random samples were normalized by coverage. For each genomic position in the hematopoiesis regulatory regions, a mean and standard deviation of coverage was calculated across the 1000 denoised random samples.

#### Identifying regions with significant changes in chromatin accessibility

For each subsample of lineage-primed HSCs, a z-score for each genomic position in the hematopoiesis regulatory regions was generated based on the normalized coverage relative to the mean and standard deviation in the 1000 denoised random samples. Regulatory “peaks” were called by combining all genomic positions with an absolute z-score greater than 2 within 200 bp of each other. The top z-scores for each peak were converted to p-values and then corrected for multiple hypothesis testing using the Benjamini Hochberg procedure. All peaks with a false discovery rate less than 0.05 were saved in a BED file. Peaks were then filtered by a minimum coverage value to remove low-coverage regions that would not be identified through typical ATAC-seq analysis.

### Data visualization

Unless otherwise specified, the WashU epigenome browser (http://epigenomegateway.wustl.edu/browser/) was for ATAC-seq signal track visualization. The denoised lineage-priming HSC subsamples (Fig. 2f) were visualized using the Integrative Genomics Viewer^33^.

### Code availability

AtacWorks is available at https://github.com/clara-genomics/AtacWorks. Custom scripts used to batch process samples for input and identify differentially-accessible regulatory regions in lineage-primed hematopoietic stem cells are available at https://github.com/zchiang/atacworks_analysis.

### Data availability

All of the data, trained models, and output signal tracks described in this paper are available for download at https://atacworks-paper.s3.us-east-2.amazonaws.com.

#### Bulk ATAC-seq

Bulk ATAC-seq datasets of various blood cell types are available from GEO under accession number GSE74912. From these datasets, B cells, NK cells, CD4+ and CD8+ T cells were used for model training, while erythroblasts and monocytes were used for testing. For the transcription factor footprinting model, NK cells, CD8+ T cells, and multipotent progenitors (MPPs) were used for training, while HSCs were used for testing. The bulk ATAC-seq dataset for Peyer’s Patch is available from ENCODE under experiment ENCSR017RQC.

#### Single-cell ATAC-seq

The dscATAC-seq dataset of hematopoietic stem cells generated for this study is available from GEO under accession number GSE147113 (reviewer access code: efqdgsooldqtpmb).

Other dscATAC-seq and dsciATAC-seq datasets are available from GEO under accession number GSE123581. From these datasets, CD4^+^ T cells, CD8^+^ T cells, and pre-B cells were used for model training, while monocytes were used for testing. Bead-isolated CD34^+^ cells were used for the combined UMAP projection. The scATAC-seq dataset of FACS-isolated peripheral blood mononuclear cells (PBMCs) is available from GEO under accession number GSE96772. These cells were used to infer cell type labels for CD34^+^ cells in the combined UMAP projection.

#### ChIP-seq

CTCF ChIP-seq tracks are available from ENCODE under experiments ENCSR000DLK (HSCs), ENCSR000ATN (B cells) and ENCSR000AUV (monocytes). H3K27ac ChIP-seq tracks are available from ENCODE under experiments ENCSR000AUP (B cells) and ENCSR000ASJ (monocytes).

## Acknowledgments

We thank Eric Xu, Joyjit Daw, and Neha Tadimeti for contributing to the code for AtacWorks. We thank Ronald Lebofsky and Giulia Schiroli for assistance in generating dscATAC-seq data. We thank Yan Hu for critical reading of the manuscript. We thank members of the Buenrostro lab and NVIDIA team for insightful comments throughout the development of this work. J.D.B., Z.D.C., and F.M.D. acknowledge support by the Allen Distinguished Investigator Program through the Paul G. Allen Frontiers Group. This work was further supported by the Chan Zuckerberg Initiative and the NIH Director’s New Innovator award. Z.D.C. is supported by the NSF-Simons Center for Mathematical and Statistical Analysis of Biology at Harvard (#1764269).

## Author Contributions

N.Y. and A.L. developed the deep learning model. A.L. and Z.D.C. performed data analysis. F.M.D. performed HSC dscATAC-seq experiments. A.L., Z.D.C., and J.D.B. wrote the manuscript with input from all authors. J.I. and J.D.B. jointly conceptualized and supervised this work.

## Supplementary Materials

Supplementary Table 1: Performance of AtacWorks on bulk ATAC-seq data from human erythroblasts. Resnet models were trained on bulk ATAC-seq data from CD4+ T cells, CD8+ T cells, B cells, and NK cells. Metrics were calculated separately on the whole genome, on chromosome 10 (not used for training), and on differential peaks (peaks present only in either the training data or the test data).

Supplementary Table 2: Comparison of AtacWorks and linear regression models on bulk ATAC-seq data from erythroblasts. The Resnet models in Supplementary Table 1 are compared against linear regression models for denoising, trained on the same data.

Supplementary Table 3: Performance of AtacWorks on bulk ATAC-seq data from erythroblasts using different sets of training data. Resnet models trained on bulk ATAC-seq data from 4 cell types (Supplementary Table 1) are compared against Resnet models trained on bulk ATAC-seq data from 1 cell type (CD4+ T cells).

Supplementary Table 4: Performance of AtacWorks on ENCODE bulk ATAC-seq data from the human Peyer’s Patch. The Resnet models are the same as in Supplementary Table 1.

Supplementary Table 5: Comparison of AtacWorks and U-Net architecture on bulk ATAC-seq data from erythroblasts. Resnet models are compared against a previously-described U-Net model architecture ^11^. All models were trained on bulk ATAC-seq data from CD4+ T cells.

Supplementary Table 6: Performance of AtacWorks on droplet single-cell ATAC-seq (dscATAC) data from natural killer (NK) cells. Resnet models were trained on dscATAC data from B cells and monocytes.

Supplementary Table 7: Performance of AtacWorks on droplet-based single-cell combinatorial indexing ATAC-seq (dsciATAC) data from monocytes. Resnet models were trained on dsciATAC data from CD4+ T cells, CD8+ T cells, and pre-B cells.

Supplementary Table 8: Breakdown of how monocyte and erythroblast ATAC-seq tracks were aggregated across donors and replicates to create paired high and low quality training data.

Supplementary Table 9: Performance of AtacWorks on low-quality bulk ATAC-seq signal from erythroblasts. A resnet model was trained on paired low- and high-quality ATAC-seq data from monocytes.

Supplementary Table 10: Performance of AtacWorks at CTCF sites in high-resolution bulk ATAC-seq data from hematopoietic stem cells (HSCs). Resnet models were trained using high-resolution bulk ATAC-seq data from MPP cells, CD8+ T cells, NK cells. Both the training and test set were limited to 200-bp regions surrounding CTCF motifs.

Supplementary Table 11: Performance of AtacWorks in predicting CTCF and H3K27ac ChIP-seq from noisy ATAC-seq. Resnet models were trained using aggregate single-cell ATAC-seq (dscATAC) data from B cells and tested on aggregate dscATAC data from monocytes.

Supplementary Table 12: Genomic regulatory regions with significant changes in chromatin accessibility relative to a permuted background in subsamples of lineage-primed hematopoietic stem cells.

Supplementary Table 13: Parameters used to train the AtacWorks Resnet models described in this paper.

Supplementary Figure 1: Clean (black) and noisy (blue) ATAC-seq signal tracks for the 4 ATAC-seq datasets (CD4+ T cells, CD8+ T cells, B cells and NK cells) used for training the deep learning model.

Supplementary Figure 2: Clean (black), noisy (blue) and denoised (green) ATAC-seq signal tracks for bulk ATAC-seq data from Erythroid cells. Detailed views of two peaks are shown. Below the noisy (blue) signal tracks, the heatmaps show the negative log of the p-value returned by MACS2 for each position and the red bars show peak calls by MACS2 using a p-value cutoff of 10^−3^. Below the denoised (green) signal tracks, the heatmaps show the probability returned by AtacWorks (representing the probability that each position is part of a peak) and the red bars show peak calls by AtacWorks using a probability cutoff of 0.5.

Supplementary Figure 3: Clean (black), noisy (blue) and denoised (green) ATAC-seq signal tracks for ENCODE bulk ATAC-seq data from the Peyer’s Patch. Detailed views of two regions are shown. Below the noisy (blue) signal tracks, the heatmaps show the negative log of the p-value returned by MACS2 for each position and the red bars show peak calls by MACS2 using a p-value cutoff of 10^−3^. Below the denoised (green) signal tracks, the heatmaps show the probability returned by AtacWorks (representing the probability that each position is part of a peak) and the red bars show peak calls by AtacWorks using a probability cutoff of 0.5.

Supplementary Figure 4: Clean (black), noisy (blue) and denoised (green) ATAC-seq signal tracks for single-cell ATAC-seq (dsci-ATAC) data from CD4+ T cells. A detailed view of one peak is shown. Below the noisy (blue) signal track, the heatmap shows the negative log of the p-value returned by MACS2 for each position and the red bar shows peak calls by MACS2 using a p-value cutoff of 10^−3^. Below the denoised (green) signal tracks, the heatmap shows the probability returned by AtacWorks (representing the probability that each position is part of a peak) and the red bar shows peak calls by AtacWorks using a probability cutoff of 0.5.

Supplementary Figure 5: Heatmaps showing the signal at 10,000 genomic regions surrounding CTCF motifs (rows) in clean (100 million read), noisy (downsampled to 5 million reads) and denoised signals.

Supplementary Figure 6: From top to bottom: Noisy aggregate single-cell ATAC-seq (dscATAC) signal from 50 monocytes (blue). ENCODE ChIP-seq signal for CTCF in monocytes (black). CTCF ChIP-seq signal predicted by AtacWorks from the noisy ATAC-seq data (green). ENCODE ChIP-seq signal for H3K27ac in monocytes (black). H3K27ac ChIP-seq signal predicted by AtacWorks from the noisy ATAC-seq data (green). Detailed views of two regions are shown, one for CTCF ChIP-seq and one for H3K27ac ChIP-seq. Red bars below the signal tracks represent peak calls from MACS2 (for noisy ATAC-seq and ENCODE ChIP-seq) or AtacWorks (for predicted ChIP-seq).

Supplementary Figure 7: Noisy and denoised ATAC-seq signal tracks for the three aggregated subsamples from Fig. 2. Each subsample was generated by selecting a single HSC and identifying the 50 most similar cells A track for the aggregated FACS-isolated HSCs is also provided for reference. Selected regions contain genes that have been implicated in lineage priming^18,23^.

## References

1. Buenrostro, J. D., Giresi, P. G., Zaba, L. C., Chang, H. Y. & Greenleaf, W. J. Transposition of native chromatin for fast and sensitive epigenomic profiling of open chromatin, DNA-binding proteins and nucleosome position. Nat. Methods 10, 1213–1218 (2013).

2. Corces, M. R. et al. Lineage-specific and single-cell chromatin accessibility charts human hematopoiesis and leukemia evolution. Nature Genetics vol. 48 1193–1203 (2016).

3. Yoshida, H. et al. The cis-Regulatory Atlas of the Mouse Immune System. Cell 176, 897–912.e20 (2019).

4. Corces, M. R. et al. The chromatin accessibility landscape of primary human cancers. Science 362, (2018).

5. Buenrostro, J. D. et al. Single-cell chromatin accessibility reveals principles of regulatory variation. Nature 523, 486–490 (2015).

6. Corces, M. R. et al. An improved ATAC-seq protocol reduces background and enables interrogation of frozen tissues. Nature Methods vol. 14 959–962 (2017).

7. Pascual, S., Bonafonte, A. & Serrà, J. SEGAN: Speech Enhancement Generative Adversarial Network. arXiv [cs.LG] (2017).

8. Yang, C. et al. High-Resolution Image Inpainting using Multi-Scale Neural Patch Synthesis. arXiv [cs.CV] (2016).

9. Liu, G. et al. Image Inpainting for Irregular Holes Using Partial Convolutions. arXiv [cs.CV] (2018).

10. Koh, P. W., Pierson, E. & Kundaje, A. Denoising genome-wide histone ChIP-seq with convolutional neural networks. Bioinformatics 33, i225–i233 (2017).

11. Rai, V. et al. Single-cell ATAC-Seq in human pancreatic islets and deep learning upscaling of rare cells reveals cell-specific type 2 diabetes regulatory signatures. Mol Metab 32, 109–121 (2020).

12. He, K., Zhang, X., Ren, S. & Sun, J. Deep residual learning for image recognition. in Proceedings of the IEEE conference on computer vision and pattern recognition 770–778 (2016).

13. Zhang, Y. et al. Model-based analysis of ChIP-Seq (MACS). Genome Biol. 9, R137 (2008).

14. Lareau, C. A. et al. Droplet-based combinatorial indexing for massive-scale single-cell chromatin accessibility. Nat. Biotechnol. 37, 916–924 (2019).

15. Neph, S. et al. An expansive human regulatory lexicon encoded in transcription factor footprints. Nature 489, 83–90 (2012).

16. Skene, P. J. & Henikoff, S. An efficient targeted nuclease strategy for high-resolution mapping of DNA binding sites. Elife 6, (2017).

17. Yu, V. W. C. et al. Epigenetic Memory Underlies Cell-Autonomous Heterogeneous Behavior of Hematopoietic Stem Cells. Cell 168, 944–945 (2017).

18. Buenrostro, J. D. et al. Integrated Single-Cell Analysis Maps the Continuous Regulatory Landscape of Human Hematopoietic Differentiation. Cell 173, 1535–1548.e16 (2018).

19. Rodriguez-Fraticelli, A. E. et al. Clonal analysis of lineage fate in native haematopoiesis. Nature 553, 212–216 (2018).

20. Pei, W. et al. Polylox barcoding reveals haematopoietic stem cell fates realized in vivo. Nature 548, 456–460 (2017).

21. Schep, A. N., Wu, B., Buenrostro, J. D. & Greenleaf, W. J. chromVAR: inferring transcription-factor-associated accessibility from single-cell epigenomic data. Nat. Methods 14, 975–978 (2017).

22. Pliner, H. A. et al. Cicero Predicts cis-Regulatory DNA Interactions from Single-Cell Chromatin Accessibility Data. Mol. Cell 71, 858–871.e8 (2018).

23. Weinreb, C., Rodriguez-Fraticelli, A., Camargo, F. D. & Klein, A. M. Lineage tracing on transcriptional landscapes links state to fate during differentiation. Science 367, (2020).

24. Li, H. et al. The Sequence Alignment/Map format and SAMtools. Bioinformatics 25, 2078–2079 (2009).

25. Quinlan, A. R. & Hall, I. M. BEDTools: a flexible suite of utilities for comparing genomic features. Bioinformatics 26, 841–842 (2010).

26. Adam, P. et al. Automatic differentiation in pytorch. in Proceedings of Neural Information Processing Systems (2017).

27. Kudo, Y. & Aoki, Y. Dilated convolutions for image classification and object localization. in 2017 Fifteenth IAPR International Conference on Machine Vision Applications (MVA) 452–455 (2017).

28. Ioffe, S. & Szegedy, C. Batch Normalization: Accelerating Deep Network Training by Reducing Internal Covariate Shift. arXiv [cs.LG] (2015).

29. Ronneberger, O., Fischer, P. & Brox, T. U-Net: Convolutional Networks for Biomedical Image Segmentation. Lecture Notes in Computer Science 234–241 (2015) doi: 10.1007/978-3-319-24574-4_28.

30. Kingma, D. P. & Ba, J. Adam: A Method for Stochastic Optimization. arXiv [cs.LG] (2014).

31. Smith, T., Heger, A. & Sudbery, I. UMI-tools: modeling sequencing errors in Unique Molecular Identifiers to improve quantification accuracy. Genome Res. 27, 491–499 (2017).

32. Li, H. & Durbin, R. Fast and accurate short read alignment with Burrows-Wheeler transform. Bioinformatics 25, 1754–1760 (2009).

33. Robinson, J. T. et al. Integrative genomics viewer. Nat. Biotechnol. 29, 24–26 (2011).

