## Supplementary figures and images for "AtacWorks: A deep convolutional neural network toolkit for epigenomics"

### Supplementary Figure 1

Clean ATAC-Seq (50 million reads)

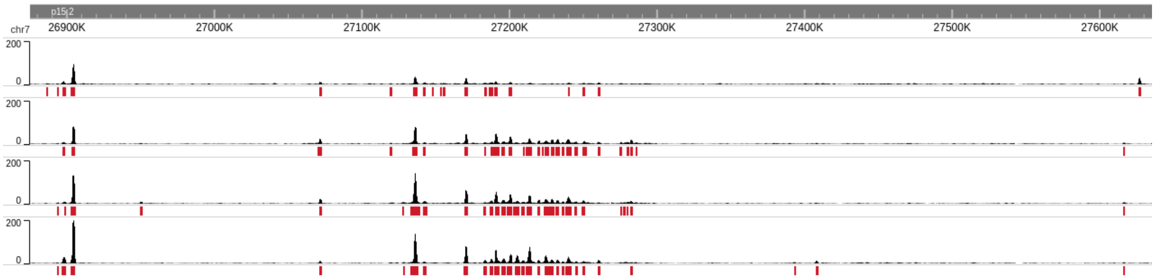


Noisy ATAC-Seq (1 million reads)

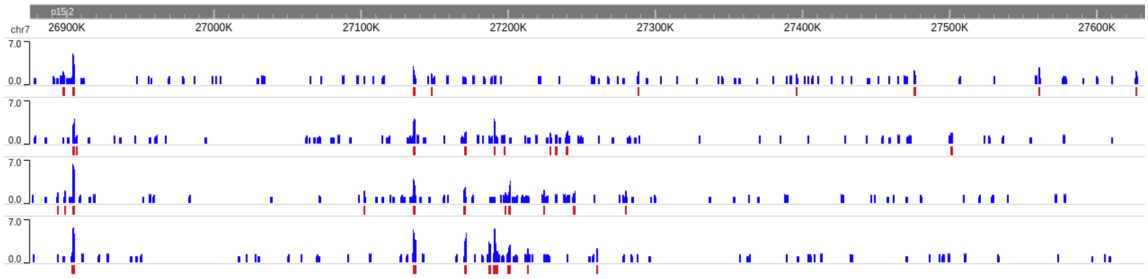

### Supplementary Figure 2

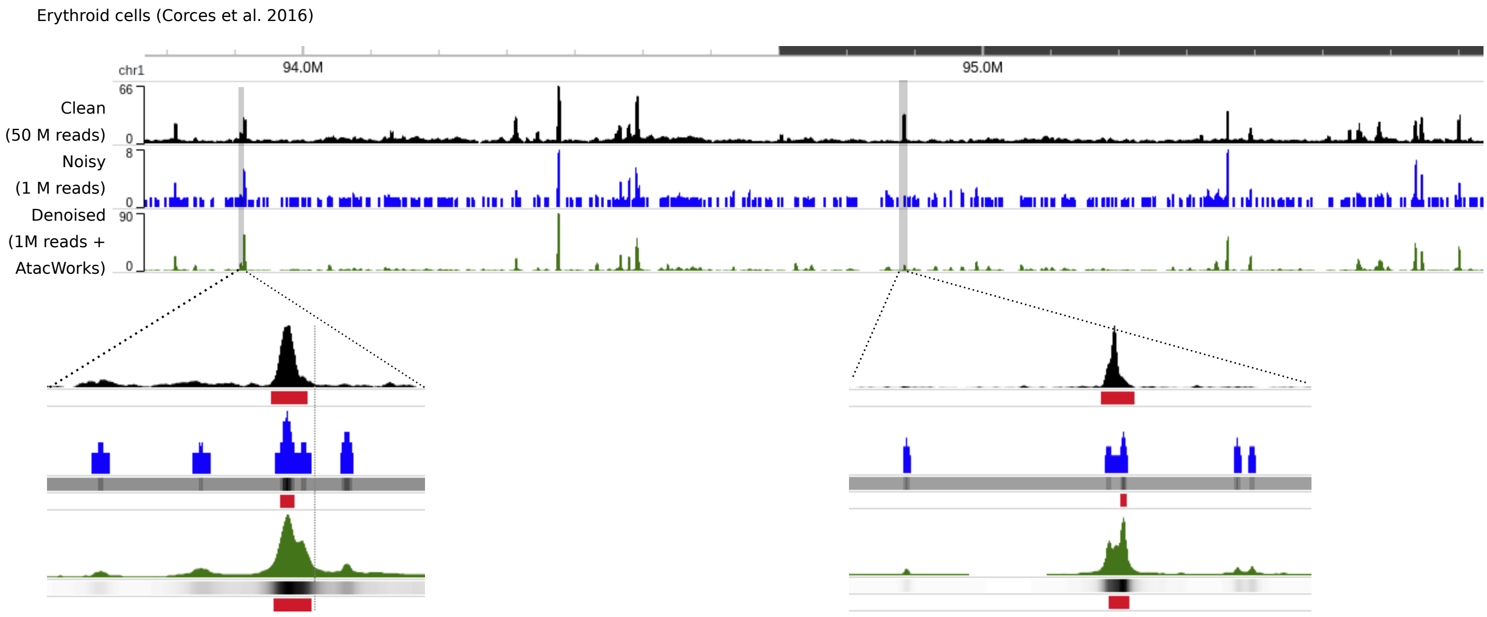

### Supplementary Figure 3

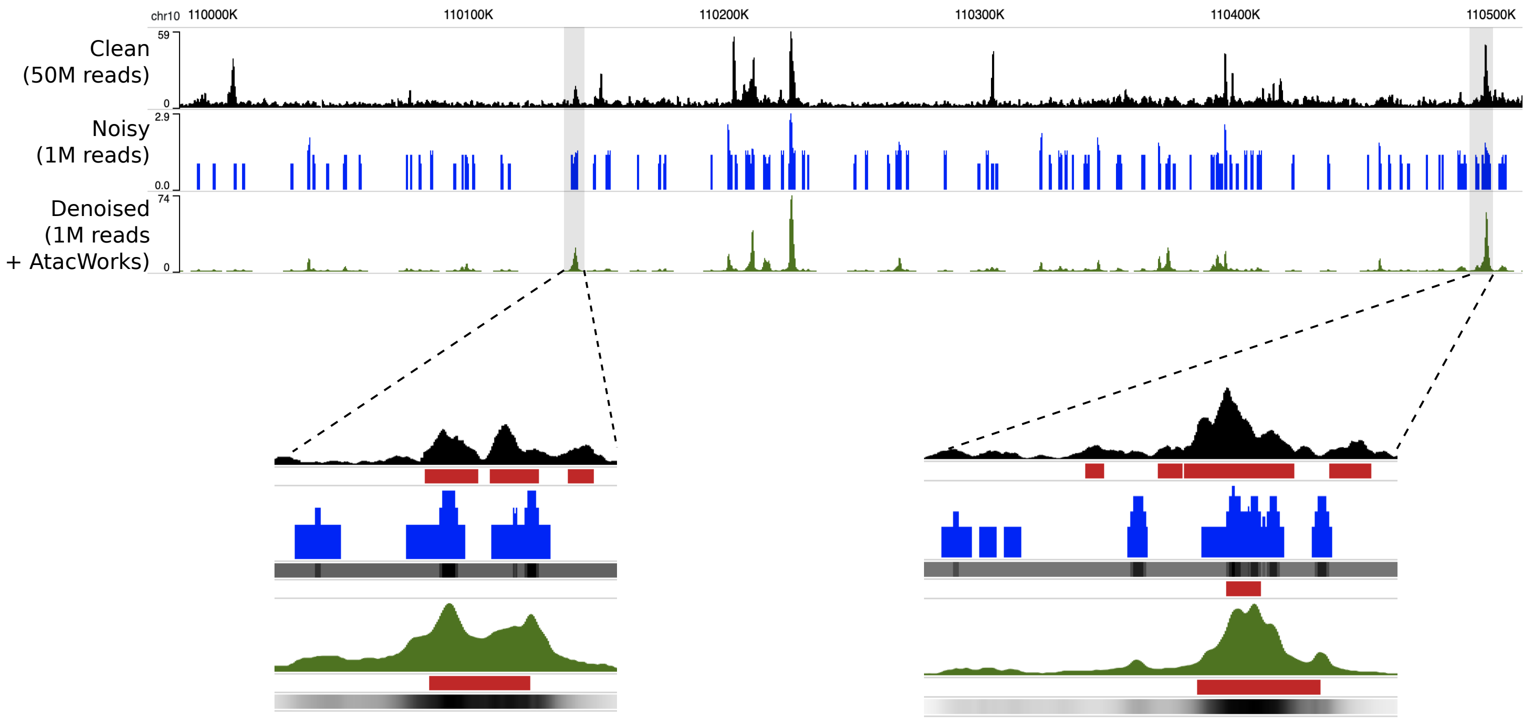

### Supplementary Figure 4

single-cell ATAC-Seq (sci-ATAC) on Monocytes (Lareau et al. 2019)

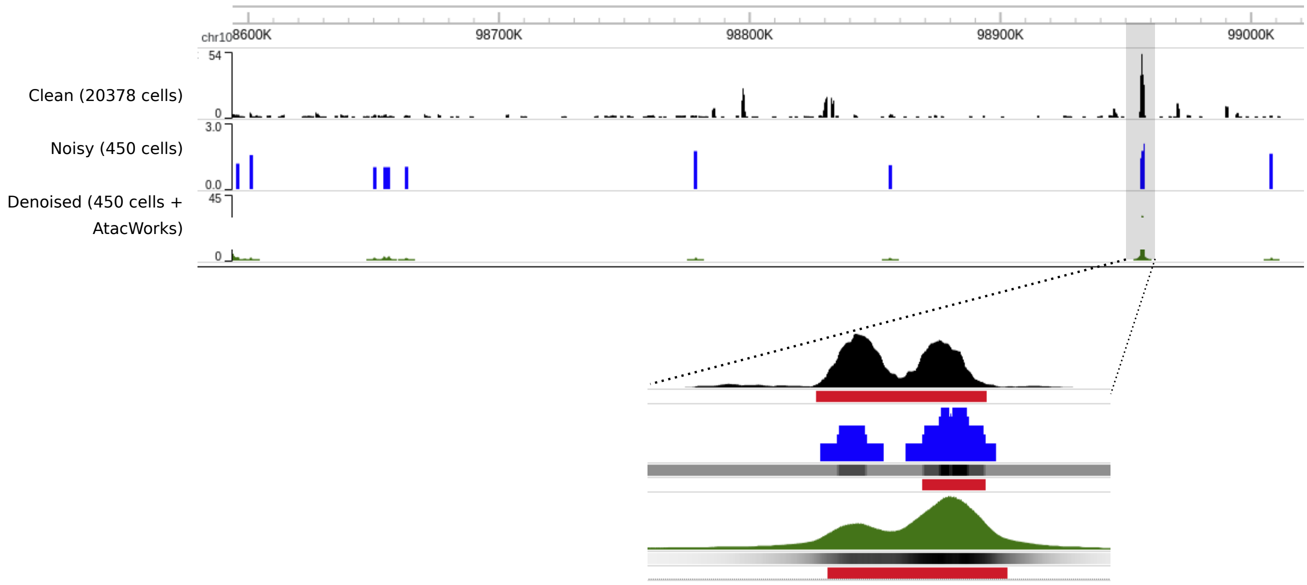

### Supplementary Figure 5

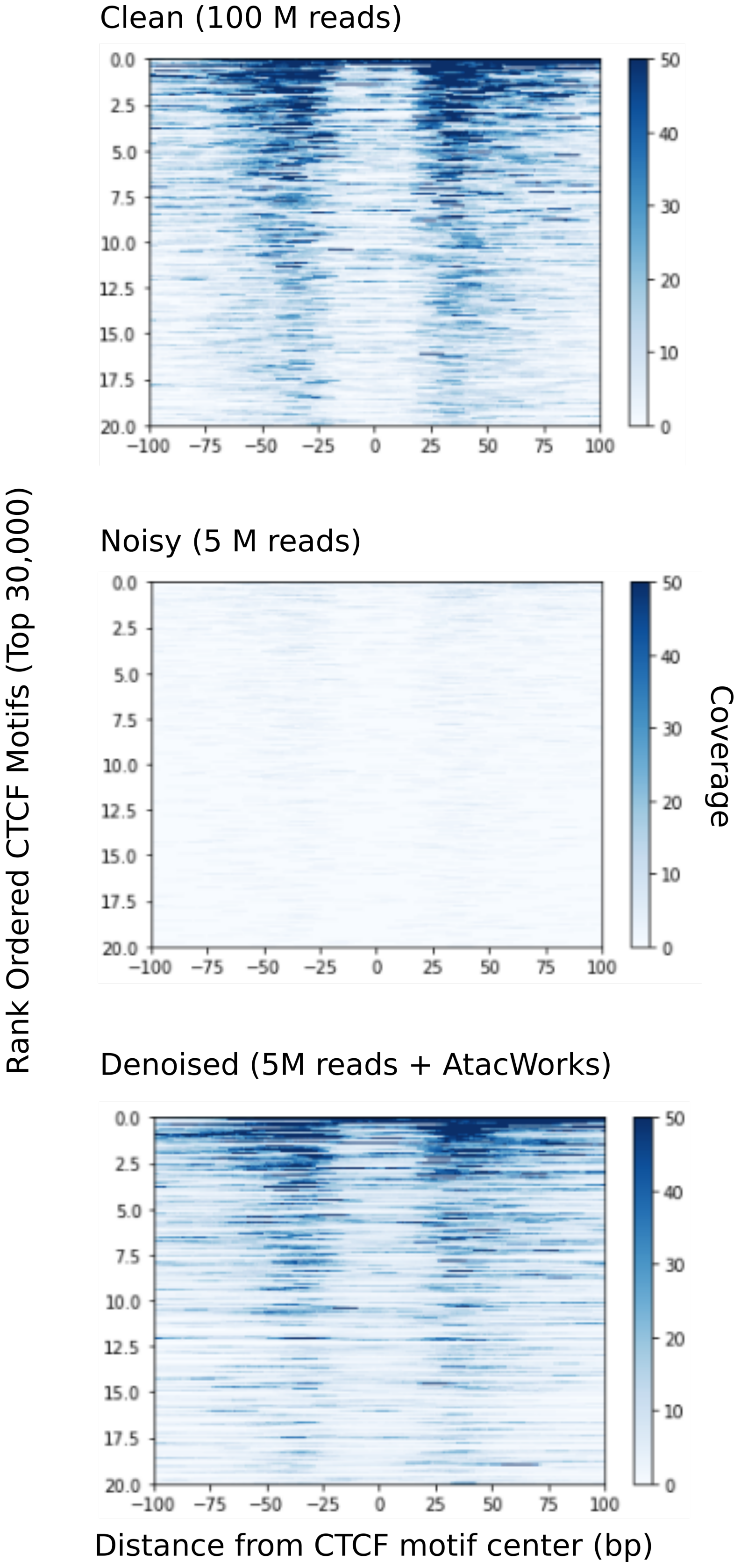

### Supplementary Figure 6

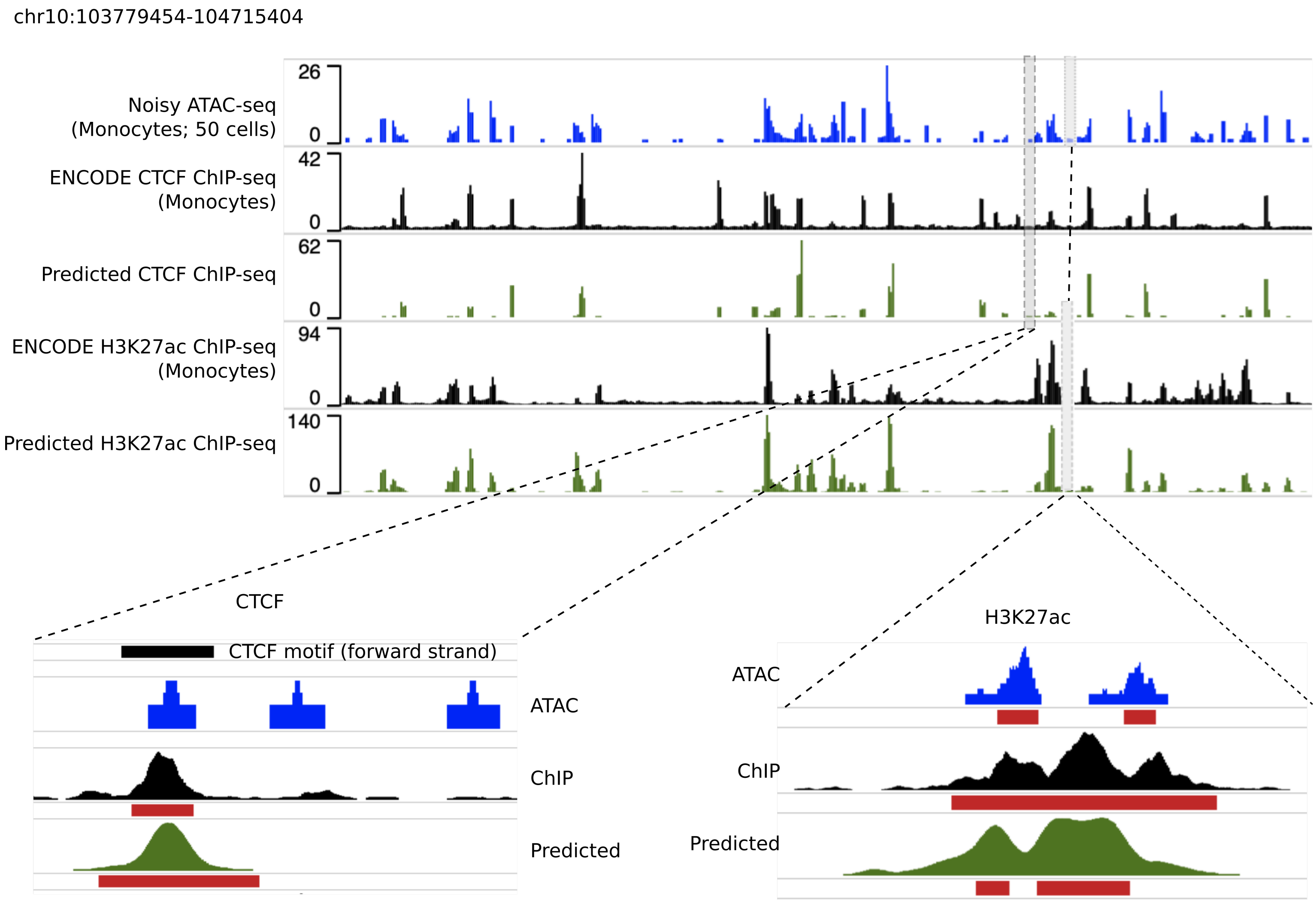

### Supplementary Figure 7

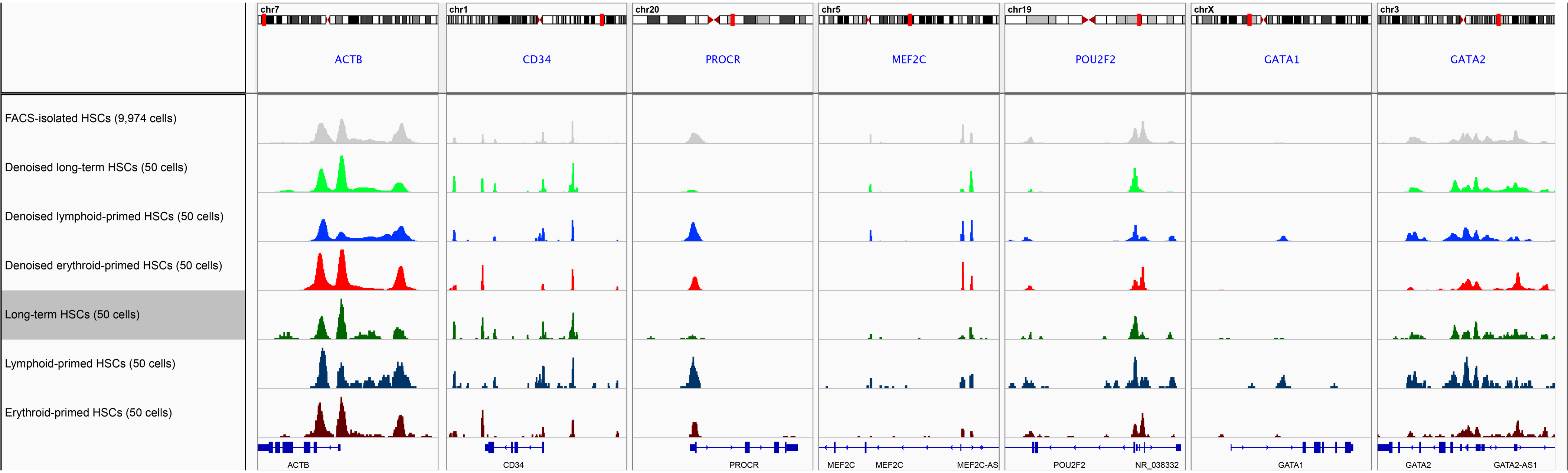
