## Supplementary Note 1 for "AtacWorks: A deep convolutional neural network toolkit for epigenomics"

**Comparison of AtacWorks with a U-Net model for ATAC-seq (PillowNet)**

Another study[1] recently reported the use of a deep learning model for denoising and peak calling from low-input ATAC-seq. Here we evaluate both AtacWorks and PillowNet (https://github.com/daquang/PillowNet), the model described in this study.

A bulk ATAC-Seq dataset from CD4+ T cells was sampled to a depth of 50 million reads to generate clean, high-coverage data, and then subsampled to a depth of 1 million reads. Both AtacWorks and PillowNet models were trained, with their respective default parameters, to learn a mapping between the noisy and clean datasets. Only a 12 Mb region of chromosome 1 was used for training and models were trained for 25 epochs.

Since the U-Net model in PillowNet predicts one output at a time, we trained two separate models, one for denoising and one for peak calling. PillowNet is set up to run on CPUs, whereas AtacWorks runs only on GPUs. The high parallelism of GPUs allows for significantly faster training and prediction.

For this small training dataset, the total time required to train both PillowNet models was 85 minutes (39 minutes for the denoising model and 46 minutes for the peak classification model) with 64 CPU cores. On the other hand, we were able to use AtacWorks to train a single model performing both denoising and peak calling in 6 minutes using a single NVIDIA V100 GPU and in 4.5 minutes using 8 V100 GPUs on an NVIDIA DGX-1 server.

We were unable to train PillowNet on a larger dataset using the provided code. Instead, we re-implemented the U-Net architecture used in PillowNet in the AtacWorks framework. We were able to train models for denoising and peak calling on the aforementioned CD4+ T cell dataset with the U-Net architecture, using the loss functions and learning rate described[1]. We also trained a standard AtacWorks resnet model to perform both denoising and peak calling using default AtacWorks parameters (Supplementary Table 13). We then applied the trained models to an ATAC-seq dataset from erythroblasts sampled to the same read depth.

We found that while the U-Net architecture performs well at both denoising and peak calling from this low-coverage dataset, the Resnet model performs better on all metrics (Supplementary Table 5).

We also note that PillowNet, as well as another previous method, Coda[2], train models that perform either denoising or peak calling, so that if both a denoised signal track and peak calls are needed, the user must train two independent models. The outputs of these models need not be correlated as the mappings they learn are independent of each other. On the other hand, AtacWorks performs denoising and peak calling jointly, so that the peak calls produced by AtacWorks from a noisy ATAC-seq dataset are a direct function of the denoised signal that it also produces.

References


1. Rai, V. *et al.* Single-cell ATAC-Seq in human pancreatic islets and deep learning upscaling of rare cells reveals cell-specific type 2 diabetes regulatory signatures. *Mol Metab* **32**, 109–121 (2020).
2. Koh, P. W., Pierson, E. & Kundaje, A. Denoising genome-wide histone ChIP-seq with convolutional neural networks. *Bioinformatics* **33**, i225–i233 (2017).
